# Printable microscale interfaces for long-term peripheral nerve mapping and precision control

**DOI:** 10.1101/688218

**Authors:** Timothy M. Otchy, Christos Michas, Blaire Lee, Krithi Gopalan, Jeremy Gleick, Dawit Semu, Louis Darkwa, Bradley J. Holinski, Daniel J. Chew, Alice E. White, Timothy J. Gardner

**Affiliations:** Department of Biology, Boston University; Neurophotonics Center, Boston University; Center for Systems Neuroscience, Boston University; Department of Biomedical Engineering, Boston University; Bioelectronics Division, GlaxoSmithKline; Department of Mechanical Engineering, Boston University; Now: Knight Campus, University of Oregon

## Abstract

The nascent field of bioelectronic medicine seeks to decode and modulate peripheral nervous system signals to obtain therapeutic control of targeted end organs and effectors. Current approaches rely heavily on electrode-based devices, but size scalability, material and microfabrication challenges, limited surgical accessibility, and the biomechanically dynamic implantation environment are significant impediments to developing and deploying advanced peripheral interfacing technologies. Here, we present a microscale implantable device – the nanoclip – for chronic interfacing with fine peripheral nerves in small animal models that begins to meet these constraints. We demonstrate the capability to make stable, high-resolution recordings of behaviorally-linked nerve activity over multi-week timescales. In addition, we show that multi-channel, current-steering-based stimulation can achieve a high degree of functionally-relevant modulatory specificity within the small scale of the device. These results highlight the potential of new microscale design and fabrication techniques for the realization of viable implantable devices for long-term peripheral interfacing.

## INTRODUCTION

There is growing evidence of the therapeutic benefit of targeted modulation of electrical signaling within the peripheral nervous system (PNS), the network of nerves and ganglia that innervate the organs and tissues of the body (Birmingham et al., 2014; Famm et al., 2013). Positive clinical trials are reported for vagal nerve stimulation to treat inflammatory diseases (Borovikova et al., 2000), depression (Sackeim et al., 2001), and epilepsy (Morris et al., 2013), and beyond these early studies, it is thought that a broad range of chronic diseases may be treatable through precise, artificial control of the PNS (Famm et al., 2013). Despite the potential of these bioelectronic therapies, progress – both in our understanding of PNS function and in developing effective modulatory “prescriptions” – will require new tools capable of making a high-quality, chronically stable interface with fine peripheral structures. A recently proposed roadmap for the development of such therapies highlights the need for new electrode-based devices that can map peripheral physiological signals and be used to precisely control end organs (Birmingham et al., 2014). Key features of these emerging interfacing technologies include high-resolution, stable recordings in single nerves, flexible nerve stimulation options that can be tailored to the physiological effect desired, and scalable platforms that can interface with small nerves of 200 µm diameter or less.

A variety of peripheral neural interfacing approaches have been previously reported – including extra-neural (Naples and Sweeney, 1988; Tan et al., 2014; Tyler and Durand, 2002), penetrating (Badia et al., 2011; Boretius et al., 2010; Christensen et al., 2014; Lago et al., 2007; Wark et al., 2013, 2014), and regenerative electrodes (Lacour et al., 2008, 2009) – each with unique sets of advantages and disadvantages for specific applications. For example, traditional extra-neural cuff electrodes interface with intact whole nerves, but have limited recording capability and are a challenge to size properly for fine nerve targets (Loeb and Peck, 1996; Navarro et al., 2005; Somann et al., 2017). Intrafascicular penetrating microelectrodes can realize significant improvements in recording and stimulating specificity, but risk substantial insertion trauma and existing platforms are too large for chronic use in many fine nerves. Regenerative electrodes can interface stably at chronic timescales, but typically entail significant nerve damage during implant and a prolonged period of compromised function during regrowth (Lacour et al., 2009; Mannard et al., 1974; Stieglitz et al., 1997). More generally, factors contributing to chronic instability and poor performance of implanted peripheral probes include mismatch in scale of device to the nerve, excessive tethering and compression forces, limited surgical access to many implant sites, and challenges of manufacturing small devices. As such, creating an implantable nerve interface targeting the fine structures of the PNS that is capable of selective stimulation and high-quality recording over chronic timescales remains an open challenge (Birmingham et al., 2014; Famm et al., 2013; Grill et al., 2009; Navarro et al., 2005; Rutten, 2002).

We previously introduced the nanoclip to harness microscale 3D printing to create interfaces with geometries tailored with near micron-resolution to the implant target, and showed that such a device could interface acutely with a small peripheral nerve (Lissandrello et al., 2017). Here, we present a nanoclip interface integrating a multi-channel thin-film electrode suitable for chronic interfacing with peripheral nerves in small animal models (Figure 1), and we demonstrate stable, longitudinal, in vivo recording of behaviorally-linked nerve activity with high signal-to-noise ratio (SNR). Furthermore, we demonstrate flexible, precision control of an end organ (here: the songbird syrinx, a complex vocal organ containing six muscles (Goller and Suthers, 1996)) to produce distinct, physiologically relevant articulatory states – in this case stereotyped, fictive vocalizations. Such stable mapping and modulation may enable longitudinal rather than cross-sectional experiments, allowing studies of how PNS dynamics are shaped by developmental processes, how changes in nerve signaling correlate with biomarker patterns during disease progression, and whether peripheral neural plasticity can be directed therapeutically via modulatory interventions (Bouton, 2017; Famm et al., 2013; Ganguly et al., 2011; Grill et al., 2009; Lütcke et al., 2013; Marder and Goaillard, 2006).

**Figure 1.**
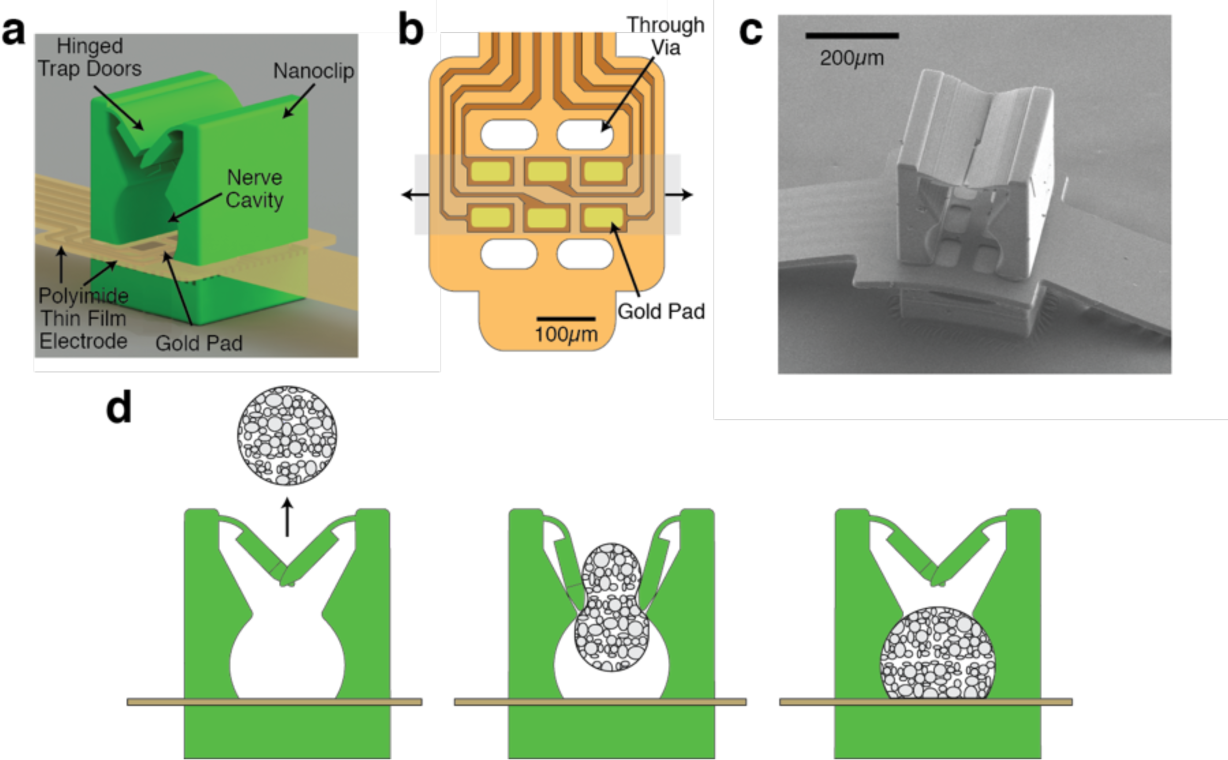
Thin-film-integrated nanoclip nerve interface overview. **(a)** Rendering of a thin-film-integrated nanoclip nerve interface showing key mechanical design components. The nanoclip was fabricated directly on the thin-film electrode via rDLW (Pearre et al., 2018) and consists of dual elastically-deformable trap doors, a nerve retention cavity, and a rigid base. **(b)** Diagram of the polyimide thin-film electrode array used in all experiments. Six gold electrode pads (45×80 µm each) were located inside the cavity (in **a**) such that they were in contact with the epineurium of the retained nerve. The thin-film through-vias allow for the mechanical integration of the nanoclip top and base with mortise-and-tenon-type joints (see Figure S2). The shaded gray region shows the location of a retained nerve; the arrows indicate the nerve axis. **(c)** SEM micrograph of the nanoclip nerve interface. **(d)** Schematic of implantation process: (left) the nanoclip is advanced towards an isolated nerve; (center) the nanoclip contacts the nerve resulting in elastic deformation of the trap doors and subsequent entry of the nerve into the central retention cavity; (right) as the nerve fills the cavity and makes contact with the electrode pads, the doors close behind the nerve to secure the nanoclip on the nerve.

## RESULTS

### Peripheral nerve interface design and fabrication

Our approach combines thin-film microfabrication photolithography techniques (Feeney and Kounaves, 2000; Madou, 2011) and high-resolution 3D printing (Sun and Kawata, 2006; Vaezi et al., 2013) to realize an implantable device that more precisely matches the scale of small nerve implant targets (Figure 1a). The thin-film electrode array is comprised of a 50nm-thick layer of gold, defining the electrode pads and interconnects, encapsulated between insulating and biocompatible polyimide layers (Lago et al., 2007) (Figure S1a). The total film thickness (12 µm) and narrow width (250 µm) of the device yield a low bending stiffness of ∼0.5 nN-m (Chang et al., 2008), which is comparable to the stiffness of peripheral tissues (Grill et al., 2009; Minev et al., 2015; Restaino et al., 2014) and correlates with a low immune response (Maiti et al., 2007; Salatino et al., 2017; Ward, 2008).

The electrode array used in the reported experiments consists of 6 gold pads (each 45×80 µm arranged in a 2-by-3 grid; Figure 1b, S2a) that make intimate contact with the extra-neural surface of the nerve (i.e., the epineurium) and are addressed individually by gold interconnects that terminate at input-output (I-O) pads at the opposite end (Figure S2a). The I-O pads were connectorized (to interface with external instrumentation) and encapsulated in polydimethylsiloxane (PDMS) to provide mechanically support and insulation (Figure S1b). The total length of the thin-film device (from electrode array to I-O pads) was 40 mm – sufficient to reach many peripheral implant targets in preclinical models while externalizing the connector at the head or back – though all six electrodes are contained within a 260 × 105 µm region at the probe head (Figure 1b, S2a). We note that the dimensions of the electrode presented here are limited only by the choice to start with commercially available photolithography techniques; microfabrication through more advanced processes could be used to produce electrodes with finer traces and thinner encapsulation (Hanson et al., 2019).

To anchor the thin-film array to the nerve, we developed a micro-scale mechanical nerve carrier – the “nanoclip” (Lissandrello et al., 2017) – that consists of two hinged trap doors flanking the entrance of a semi-cylindrical cavity that passes through the body of the small device (∼300×300×400 µm) and securely retains the nerve against the electrodes (Figure 1a, S2b-d). The nanoclip is fabricated using a novel two-photon direct-write lithography technique developed for this application (Pearre et al., 2018). The approach enables the seamless integration of an acrylic photopolymer with the thin-film array (Figure 1c, S1b) without additional manual assembly steps. In addition, this fabrication method offers the ability to tailor the clip to dimensions of the targeted nerve. This ensures a fit with near-micron precision (Figure 1c), limited only by pre-surgical measurement of nerve diameter. Following nanoclip printing and post-processing (Figure S1b), the gold electrodes were plated with electrodeposited iridium oxide films (EIROF) (Cogan, 2008; Gillis et al., 2018), increasing their charge injection capacity (Figure S3) and reducing the likelihood of irreversible electrochemical processes at the electrode surfaces during stimulation (Cogan et al., 2016; Meyer et al., 2001).

### Uncompromised nerve function following implant

Recent studies suggest that some approaches to PNS interfacing are associated with significant compromise of nerve health at both acute and chronic timepoints (Rydevik et al., 1981; Somann et al., 2017; Tyler and Durand, 2003), casting doubt on prior findings establishing the basic physiology of the periphery and the feasibility of bioelectronic therapeutic and restorative technologies. Thus, one of the principal design constraints was to develop an interface that minimized surgical complexity such that insult to the targeted nerve and the surrounding tissues would be minimal. For implantation, the nanoclip is advanced towards the nerve (Figure 1d, left), resulting in elastic deformation of the trap doors (Figure 1d, center) and subsequent entry of the nerve into the central retention channel. After the nerve clears the doors, they return to their original configuration, securing the nerve inside the retention channel (Figure 1d, right). The nanoclip is sized for a snug, but not tight fit, with complete closure of the hinges and minimal radial compression of the nerve (see Figure S2b-d for device dimensions). With this design and fit, all devices remained on the nerve for the entirety of the acute and chronic tests reported.

To evaluate the chronic safety of the design, we implanted the nanoclip on the songbird tracheosyringeal nerve (nXIIts) – an avian hypoglossal analogue that innervates the songbird vocal organ, the syrinx, (Figure 2a) and shows strong homologies to mammalian sensorimotor nerves, containing approximately 1000 myelinated and unmyelinated fibers (Lissandrello et al., 2017; Wild, 1997) – reasoning that any disruption of nerve function would be revealed in acoustic distortion of the otherwise highly-stereotyped song (Lissandrello et al., 2017; Ölveczky et al., 2011). We recorded song from adult male zebra finches (n = 3 birds) for three days before and eight days after bilaterally placing nanoclips on the nXIIts. In all experiments, birds resumed singing the day after surgery, robustly producing well-structured vocalizations from the first utterances. To characterize changes in nerve function following implant, we quantified the moment-by-moment spectral similarity of pre-implantation song to song produced in subsequent days (see Methods). We found a small but not significant difference in song acoustic structure produced before and after implant – consistent with spectral changes measured in birds receiving bilateral sham implants (n = 3) and significantly less than in birds receiving bilateral nerve crush injury (n = 3) (Figure 2b).

**Figure 2.**
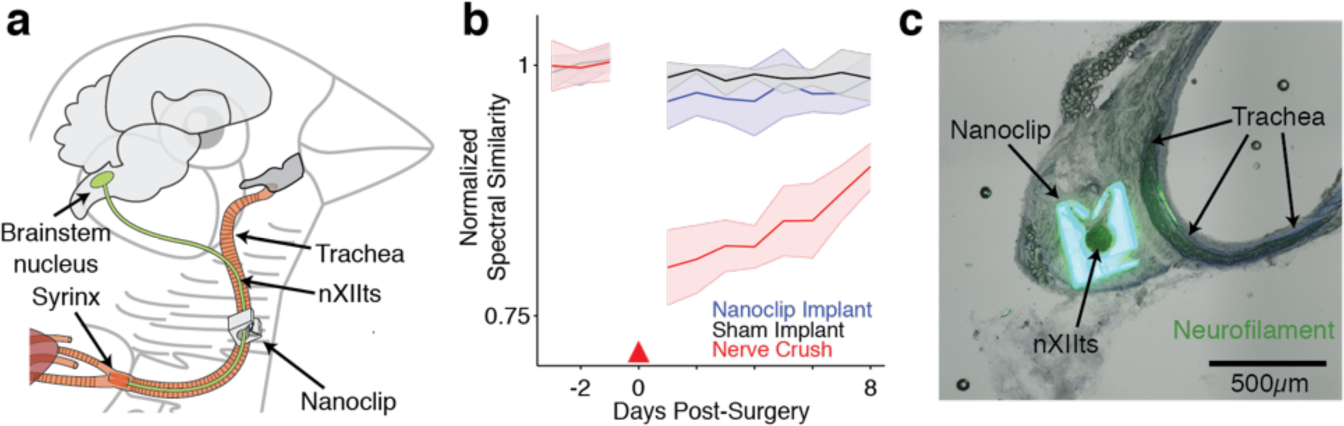
Nanoclip nerve interface implantation and bio-integration. **(a)** Diagram of the nanoclip interface implanted on the nXIIts nerve of the zebra finch. **(b)** Functional recovery of nXIIts following bilateral nanoclip implant (blue), sham implant (black), and nerve crush (red) as assessed by comparing pre- and post-manipulation song acoustic structure (see Methods). Line and shaded region denote mean +/-std across n=3 birds per condition. Red triangle marks day of surgery. **(c)** Micrograph of a cross-section of the nanoclip, nXIIts nerve, and host tissue 11 months post-implant shows that the device remains centered on the nerve with minimal penetration of fibrotic tissue into the retention cavity despite complete encapsulation of the device. Antibodies against neurofilament shown in green; color of the nanoclip is due to photoresist auto-fluorescence.

In chronically implanted birds for which post-mortem histological sectioning was performed (n = 4), implanted nanoclips were found to be securely latched over the nerve, with minimal qualitative evidence of host tissue insult. Histological inspection showed the nanoclip implant provokes a typical fibrotic response, completely encapsulating the device within weeks. Interestingly, we found limited fibrotic penetration between the nerve and nanoclip up to 11 months after implant (Figure 2c), suggesting the potential for long-term viability of the preparation (Caravaca et al., 2017; Hiebl et al., 2010; Xiang et al., 2016; Xue et al., 2015).

### High-SNR acute recordings from fine peripheral nerves

To validate the nanoclip for in vivo recording of nerve activity, we recorded evoked compound responses from the nXIIts of adult male zebra finches under anesthesia. The nXIIts was exposed by blunt dissecting the connective and adipose tissue surrounding the trachea, and a ∼1 mm section of the nerve was isolated. The device was implanted on the nerve, with the electrode pads making contact with the epineurium (Figure 3a). Silver wire bipolar hook electrodes were placed on a similarly isolated section of the nXIIts approximately 15-20 mm distally for bulk stimulation of nerve responses. Biphasic stimulation pulses (200 µs/phase) were applied at 1 Hz, and data were recorded from 5 ms before to 25 ms after the stimulation onset. This process could be continued indefinitely, within the limits of the anesthesia protocol.

**Figure 3.**
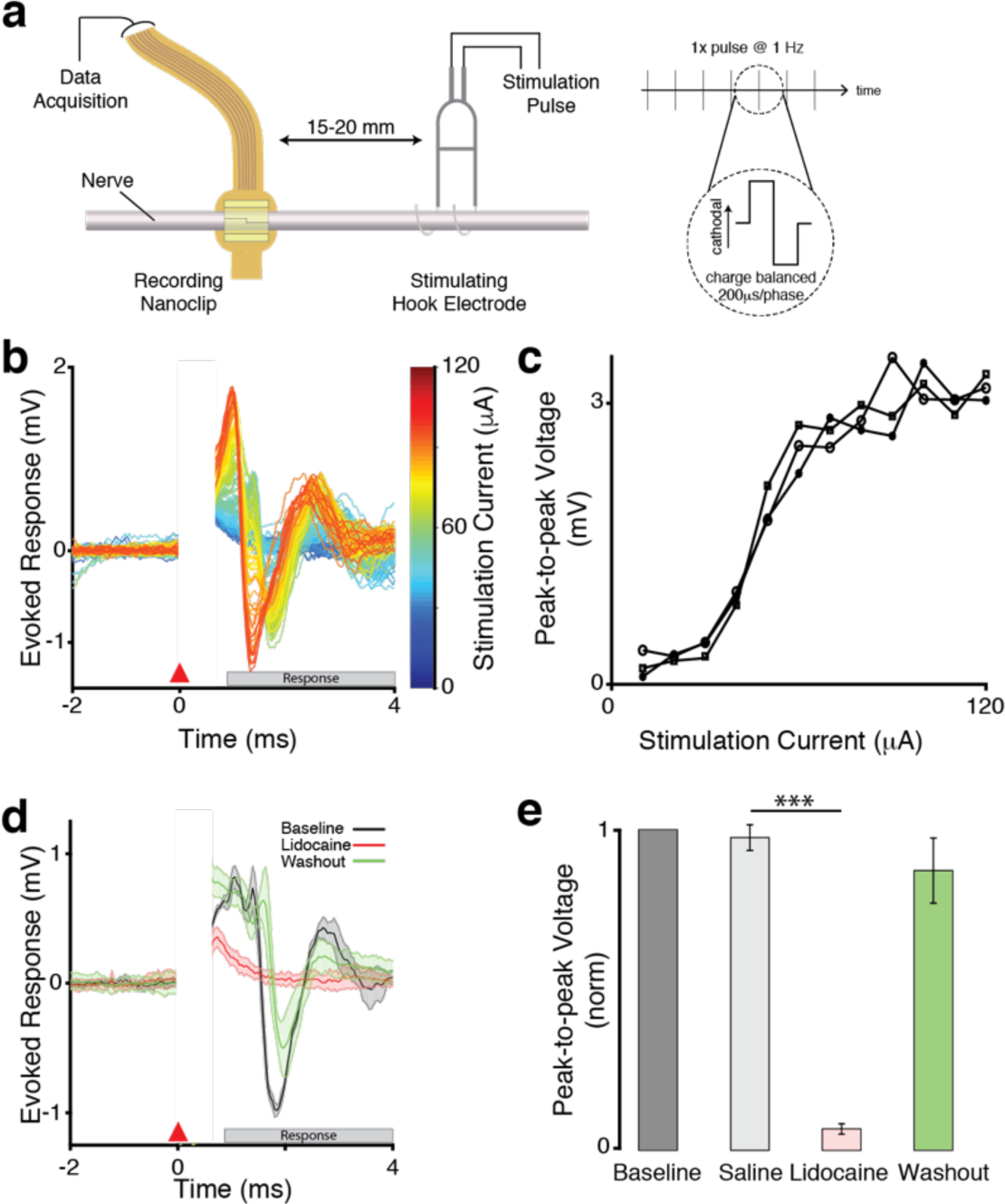
Nanoclip-anchored thin film arrays produce high-quality electroneurograms in acute preparations. **(a)** Schema for acute recording of evoked compound action potentials. (Left) Current-controlled stimulation was delivered via bipolar silver hook electrodes; evoked responses were recorded by a nanoclip interface implanted 15-20 mm rostrally. (Right) Biphasic, cathodal-leading stimulating pulses, 200 μs/phase at 10-120 μA, were delivered at 1 Hz for 16-24 trials. **(b)** Example of graded evoked response to increasing stimulation intensities. Stimulation applied at t=0 ms; stimulation artifact appears t=0-0.5 ms; evoked response follows. **(c)** Evoked response peak-to-peak voltage showed an expected sigmoidal relationship with stimulation intensity. Each data point indicates the mean across trials within an animal (n=3 birds; 16-24 trials each per stimulation intensity). **(d)** Example of stimulation evoked responses recorded before, during, and after local lidocaine application at the recording site. Each condition shows mean +/-standard error of the mean (sem) (shaded) for n=16 trials; all responses evoked at 64μA. **(e)** Evoked response peak-to-peak voltage for different experimental conditions. Response amplitudes were normalized to baseline condition to facilitate comparison across animals (n = 3 birds); error bars indicate sem. \*\*\**P* < 0.001. For tests of significance for this and all other figures see Methods.

We obtained graded evoked response curves by varying the stimulation current amplitude (n = 3 birds; Figure 3b). Consistent with estimates of nerve conduction velocities in myelinated axons of a similar size (Andrew and Part, 1972), the most salient features of the evoked responses occurred ∼0.75 ms post-stimulation and persisted for up to 4 ms. The peak-to-peak voltages of evoked responses showed a sigmoidal relationship with stimulation intensity (Figure 3c), as expected (Gruner and Mason, 1989). The minimal evoked response detected by the interface was 37.6 +/-6.5 µV with an SNR of 8.4 +/-0.95 dB; at the mean stimulation intensity assayed (i.e., 60 µA), evoked responses were recorded with an SNR of 48.1 +/-4.2 dB (see Methods). To confirm that the recorded responses were of neural origin, we blocked nerve transduction with lidocaine (2.0% in saline) applied at the stimulating site (n = 3 birds; Figure 3d); following saline washout, evoked response amplitudes were not significantly different from sham controls (Figure 3e).

### Stable, longitudinal mapping of behaviorally-linked activity in a small peripheral nerve

One of our principal goals in developing the nanoclip was to realize a device capable of making high-quality recordings of small nerve activity over an extended period of time. In adult zebra finches, the motor program underlying crystallized song is composed of highly-stereotyped patterns of cortically-generated neural activity that are precisely time-locked to the song (Fee et al., 2004). Thus, to assess the long-term stability and reliability of the interface, we chronically implanted the nanoclip onto the nXIIts, the sole source of innervation to the songbird vocal organ (Suthers and Margoliash, 2002; Wild, 1997), and recorded singing-related nerve activity from freely-behaving animals (n = 5; Figure 4a).

**Figure 4.**
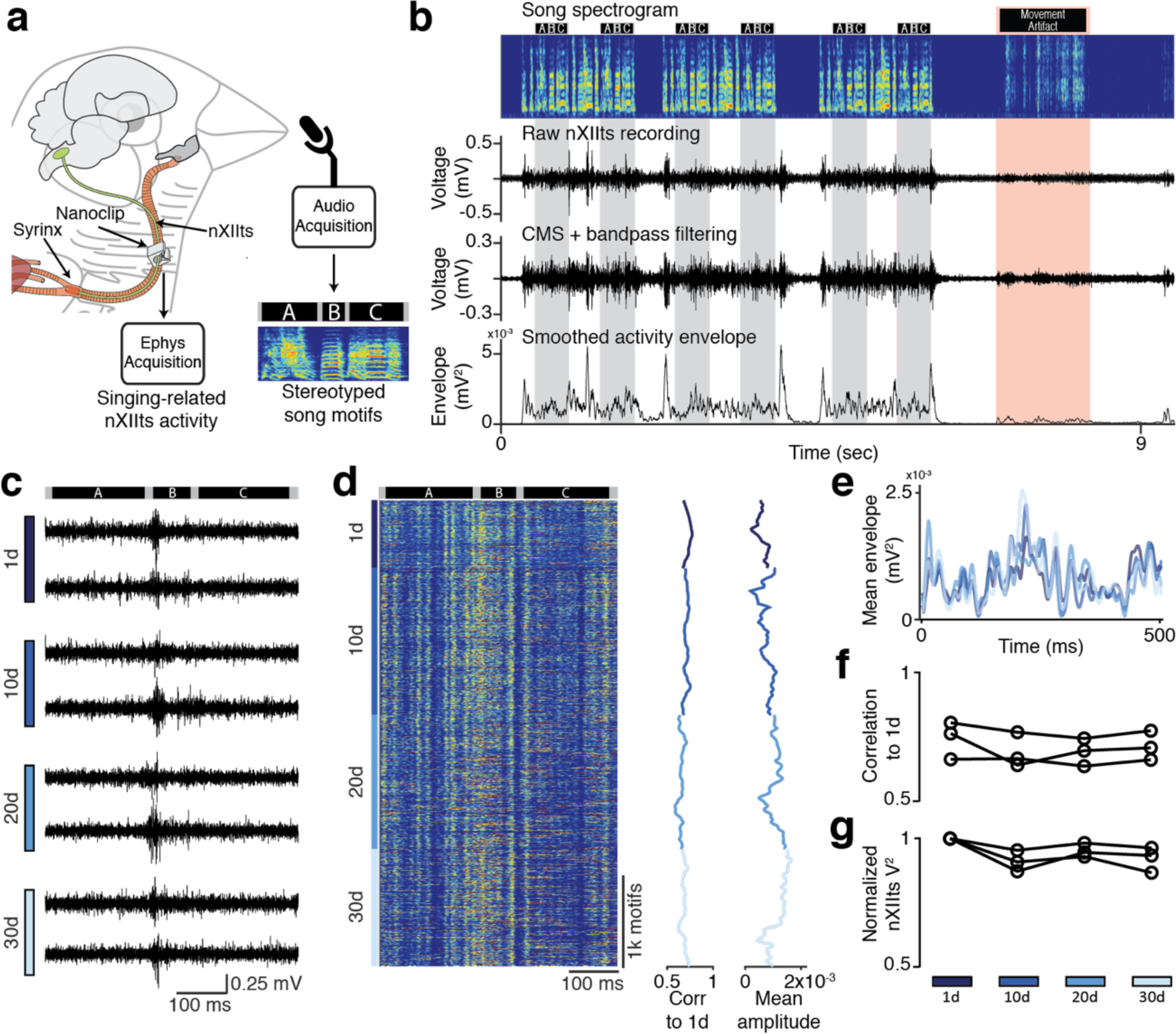
Long-term stable recordings with minimal degradation over more than four weeks. **(a)** Schema for chronic recording of nXIIts activity in the freely-behaving songbird. A nanoclip was implanted on the right-side nXIIts ∼15mm rostral to the syrinx. Song-triggered acquisition software captured both vocalization and concurrent nXIIts activity. Though all vocalizations were recorded, the principal focus of these studies were the stereotyped song motifs. Example motif spectrogram is from bird shown in b-e; black bars above the spectrogram indicate the beginning and end of each syllable in the motif (‘A’, ‘B’, and ‘C’). **(b)** Example of chronic nXIIts recording aligned to song. Top row: spectrogram of bird’s song containing three song bouts, each with two repeated motifs. Black bars above the spectrogram indicate song syllable boundaries; gray bands denote song motifs. Red band identifies movement-related noise and the associated motion artifact in the electrophysiology recording. 2^nd^ row: unprocessed nXIIts activity shows temporal correlation with song production. 3^rd^ row: nXIIts activity (from 2^nd^ row) after common-mode subtraction (CMS) and bandpass filtering. Bottom row: smoothed neural activity envelope. **(c)** Representative chronic recordings from the nXIIts nerve on days 1, 10, 20, and 30 post-implant. Dark bars at top indicate the recording alignment to syllables in the motif. **(d)** Stability of chronic recordings from bird in b, c over 30 days. Left: song-aligned nXIIts activity envelopes for motifs produced on Days 1, 10, 20, and 30 post-implant. Middle: correlation between the song-aligned nXIIts activity pattern and the average Day 1 activity pattern. Right: mean nXIIts signal envelope amplitude during singing**. (e)** Average song-aligned signal envelope (over 10 trials) recorded on days 1, 10, 20, and 30 post-implant. Colors identify recording days. **(f)** Stability over time of song-aligned nXIIts activity measured as the Pearson’s correlation to the average activity pattern on the first day of singing and averaged across n = 3 birds (Methods). Data points correspond to 25 song motifs. **(g)** Similar to (f), but showing stability over time of the mean nXIIts signal envelope amplitude during singing, normalized to the first day of singing.

As in recordings from singing-related CNS structures (Ali et al., 2013; Guitchounts et al., 2013; Ölveczky et al., 2011), we observed robust nXIIts activity that was correlated with song behavior (Figure 4b, gray bands). This correlation survived common mode subtraction, consistent with the recorded signals being of neural origin. That there were no similarly robust voltage fluctuations during periods of vigorous movement (Figure 4b, red band) additionally supports these signals as reflective of nerve activity rather than movement-related artifact. Representative recordings at 1, 10, 20, and 30 days post-implant showed well-defined signals with modulation amplitudes of ∼500 µV (Figure 4c; see Methods).

In a subset of animals (n = 3), we were able to record singing-related nXIIts activity for more than 30 days (31, 34, and 37 days). These recordings showed a remarkable degree of stereotypy in the song-aligned neural signal envelopes over the 30-day period (Figure 4d,e). Across n = 3 birds, we found no significant difference in the correlation between song-aligned nXIIts activity patterns and the mean activity pattern on day 1 (Figure 4e,f); mean signal envelope amplitudes declined modestly but not-significantly over this same period (Figure 4g). The ability to make stable longitudinal recordings of the complex nXIIts dynamics underlying song suggests that the nanoclip interface is uniquely suitable to mapping PNS function at chronic timescales.

### Precision control of a fine peripheral nerve to evoke complex fictive behavior

Precise in vivo stimulation of peripheral nerves is essential for probing the function and for providing therapeutic control of limbs and end organs (Andersson and Tracey, 2012; Birmingham et al., 2014; Raspopovic et al., 2014). Using a two-nanoclip experimental preparation, we recorded compound responses with a rostrally-placed nanoclip that were evoked with a second nanoclip interface implanted 15 mm caudally (Figure 5a). Biphasic stimulation pulses (200 µs/phase) were applied at 1 Hz, and data were recorded for 5 ms before and 25 ms after the stimulation onset. We obtained graded evoked responses by varying the stimulation current (Figure 5b). Across experiments, we found stimulation thresholds of 9.2 +/-1.6 µA (n = 3 birds), suggesting substantial isolation of the electrodes from the surrounding tissue. We note that these stimulation thresholds were consistently lower than those from bipolar silver hook stimulation electrodes (Figure 3), and indeed are within the range reported for intrafascicular stimulation (Branner and Normann, 2000; Gillis et al., 2018).

**Figure 5.**
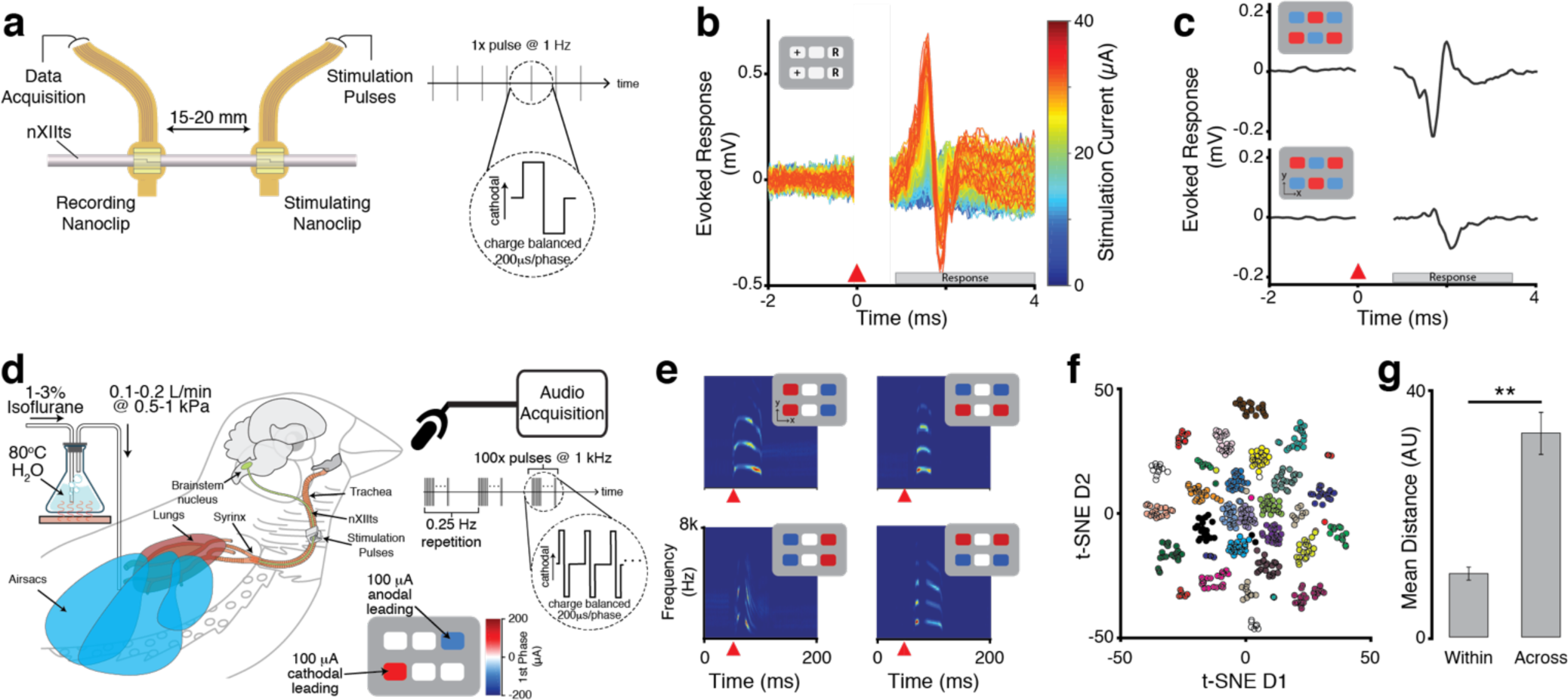
Multi-channel stimulation in a small tissue volume can achieve a high-degree of modulatory specificity. **(a)** Schema for acute multi-channel stimulation of evoked compound action potentials. (Left) Current-controlled stimulation was delivered via a rostral nanoclip interface; evoked responses were recorded by a second nanoclip interface placed 15-20 mm caudally on the same nerve. (Right) Biphasic, cathodal-leading stimulating pulses, 200 μs/phase at 5-40 μA, were delivered at 1 Hz. **(b)** Example of graded evoked response to increasing stimulation intensities. Stimulation applied at t=0 ms; stimulation artifact appears t=0-0.5 ms; evoked response follows. Inset shows stimulation electrode configuration – ‘+’ denotes source of cathodal first phase; ‘R’ denotes electrodes for the current return path. **(c)** Evoked compound responses show that spatially distinct patterns of stimulation can produce a high-degree of modulatory specificity. Inset shows the location of electrodes mediating stimulation pulses with cathodal (red) or anodal (blue) leading phases. **(d)** In vivo experimental setup for current-steering-evoked fictive singing. The nanoclip was implanted on the right-side nXIIts of an anesthetized adult male zebra finch, and an air cannula was placed in the abdominal air sac (see Methods). The respiratory system was pressurized by flowing warmed and humidified (35C; 80%) isoflurane dissolved in oxygen through the cannula (0.1-0.2 L/min at 0.5-1.5 kPa). Multi-channel stimulation (100 biphasic current pulses at 1 kHz, 200 μs/phase) with unique spatial patterns were applied to the nXIIts with the nanoclip interface, eliciting audible vocalizations that were recorded via a microphone. **(e)** Representative examples of current-steering-elicited fictive vocalizations. For each pairing, the inset (upper right of each spectrogram) identifies the stimulation pattern that produced the vocalization shown in the spectrogram. Each spectrogram depicts the vocalization from a single trial; the orange triangle marks the onset of stimulation. **(f)** Two-dimensional t-SNE embedding of high-dimensional representations of fictively-produced vocalizations (i.e., spectrograms) from one experiment testing 24 distinct current-steering stimulation patterns. Data points correspond to individual trials; colors denote current-steering patterns (see Figure S4 for pattern definitions). **(g)** Mean pairwise distance between t-SNE embedded fictive vocalizations within and across multi-channel stimulation patterns. Gray bars show mean across all tested stimulation patterns in each experiment; error bars indicate std across n = 6 birds.

Having demonstrated the capacity for bulk nerve stimulation using the nanoclip, we wondered whether precisely steering stimulation currents between the six electrodes would enable access to a wider range of nerve activation states. Using the two-nanoclip preparation (Figure 5a), we recorded compound responses evoked using a set of spatially distinct stimulation patterns (biphasic pulses, 200 µs/phase @ 1 Hz). We found that compound responses with unique waveforms and amplitudes could be evoked by changing the spatial stimulation pattern while keeping total current flow constant across trials (Figure 5c). This suggests that, despite the small scale of the nanoclip, multi-channel, current-steering-based stimulation can nevertheless achieve a modulatory specificity over that of mere bulk stimulation.

Though differences in the waveforms of the current-steering-evoked responses were visually apparent (Figure 5c), it remained unclear whether such differences in nXIIts activation were of functional significance. To test whether the nanoclip was capable of functional modulation of fine peripheral nerves, we developed a novel acute preparation – “fictive singing” – that offers complete control over the vocal-respiratory system of the anesthetized songbird. As in human speech, zebra finch vocalization requires both mechanical control of the vocal organ and elevated respiratory system pressure. Hence in the fictive singing paradigm, the muscles of the syrinx are articulated by multi-channel stimulation of the nXIIts while the respiratory system is pressurized to levels typical of normal singing (Aronov et al., 2011) by a cannula placed in the abdominal airsac (Brown and Pilny, 2006) (Figure 5d). We found that unilateral stimulation of the nXIIts (100 biphasic pulses, 200 µs/phase @ 1 kHz) was sufficient to reliably transition the vocal system from quiescence, to a sound-producing state, and back to quiescence (Figure 5e; see Methods). This demonstrates the nanoclip capable of stimulating small nerves to a physiologically efficacious state.

To functionally validate the nanoclip for precision nerve control, we quantified differences in the spectral structure of fictive vocalizations evoked by spatially distinct patterns of stimulation. We found that repeated application of a single current steering pattern in this preparation could reliably produce stereotyped fictive vocalizations with spectrotemporal characteristics consistent with species-typical song elements. In addition, we observed that spatially inverted current steering patterns often elicited fictive vocalizations that were acoustically distinct from those produced by the initial pattern (Figure 5e). To quantify differences in the acoustic structure of fictive vocalizations produced by the full set of tested stimulation patterns (Figure S4), we embedded high-dimensional representations of the fictive vocalizations (i.e., high-resolution spectrograms; see Methods) in two-dimensional space using t-distributed stochastic neighbor embedding (t-SNE) (van der Maaten and Hinton, 2008). We found that this projection of vocalization spectral structure produced plots containing clusters that well-demarcated distinct current steering patterns (Figure 5f). To quantify the reliability and specificity of current steering stimulation with the nanoclip, we compared the mean pairwise distance in the embedded space between each vocalization produced by a given stimulation pattern against the mean pairwise distance between vocalizations produced by different stimulation patterns. Across experiments (n = 6 birds), we found within-pattern distances were shorter than across-pattern distances (Figure 5g). This suggests both that functionally-overlapping subpopulations of nerve fibers could be reliably activated upon successive applications of the same current-steering pattern and that non-identical subpopulations were activated by other patterns.

## DISCUSSION

The development of new devices and approaches for the detection and modulation of signaling patterns within the nervous system puts within reach a new class of bioelectronic therapies (Birmingham et al., 2014; Famm et al., 2013). The PNS will be at the center of these treatments, as the functions it controls in chronic disease, its hierarchical organization whereby branches diverge to become more functionally specific and homogenous, and its relative surgical accessibility in comparison to intracranial structures render it tractable to targeted intervention. For these approaches to become viable, interfacing technologies that not only nominally ‘fit’ the implant target, but also match the overall scale, geometry, and biomechanics of these nerves will be necessary. However, conventional chronic nerve interfaces have geometries and mechanical properties that remain orders of magnitude divergent from those of the implant targets, increasing the risk of insult to host tissues, incidental damage to the nerve, and diminished efficacy due to poor compliance with the nerve (Grill et al., 2009; Minev et al., 2015; Salatino et al., 2017; Somann et al., 2017; Tyler and Durand, 2002).

To begin to address these challenges, we developed the nanoclip – a printable, safe microscale nerve interface (Figure 1,2) that achieved high-resolution, stable recordings from a small diameter peripheral nerve over multi-week timescales (Figure 3,4). In addition, we demonstrated that this device can achieve a high degree of modulatory specificity of fine nerves and end organ effectors (Figure 5). To our knowledge, a device capable of similarly safe, stable, and long-term chronic recording and precise modulation of peripheral structures of this size has not been presented previously.

Relating nerve signaling is to physiological function is a core aim of peripheral neuroscience and a prerequisite to the realization of bioelectronic treatments as a viable clinical therapy. This is most commonly investigated by recording or modulating peripheral signals during discrete experimental preparations, but such intermittent interventions only provide snapshots of nerve function and fail to address how function is modulated outside of the experimental context or across physiological states (Arieli et al., 1996; Dhawale et al., 2017; Evarts, 1964; Gulati et al., 2014; Lee and Dan, 2012; Santhanam et al., 2007). In addition, intermittent recordings are ill-suited for reliably tracking signals over time (Dickey et al., 2009; Fraser and Schwartz, 2012; McMahon et al., 2014; Santhanam et al., 2007; Tolias et al., 2007), making it difficult to discern how peripheral dynamics are shaped over longer timescales by developmental or disease processes (Bouton, 2017; Famm et al., 2013; Ganguly et al., 2011; Grill et al., 2009; Lütcke et al., 2013; Marder and Goaillard, 2006). Furthermore, the development of future closed-loop bioelectronic therapies that record, decipher, and modulate ongoing activity will require continuous access to high-quality, stable nerve recordings (Birmingham et al., 2014; Famm et al., 2013; Raspopovic et al., 2014; Reardon, 2014). Addressing such fundamental challenges may be greatly aided by the nanoclip’s ability to make a longitudinally stable, bi-directional interface in freely-moving animals. Such chronic preparations will allow probing of small nerve activity and function over more trials, experimental conditions, and behavioral states, thus increasing the power with which inferences about PNS function can be made (Lütcke et al., 2013).

A number of technical challenges remain. Most neural interfaces — even those composed of flexible thin-films like polyimide (here and ref (Hiebl et al., 2010; Lago et al., 2007; Seo et al., 2016)) and parylene (Caravaca et al., 2017; Chamanzar et al., 2015) — present elastic moduli substantially above the kilopascal-range of the host tissues (Mahan et al., 2015; Novak et al., 2012; Patel et al., 1998), aggravating long-term reactive responses and limiting interface longevity. High-resolution 3D printing offers the possibility of orthogonalizing the mechanical and material properties of an object, such that pliable microscale structures can be created from materials that are otherwise stiff and rigid (Bückmann et al., 2012, 2014; Lee et al., 2012). This decoupling is evident in the nanoclip’s elastically-deformable hinges (Figure 1) despite the Young’s modulus of their bulk material being within the gigapascal range (Meza et al., 2015), and additional investigation of metamaterial designs that mimic the biomechanics of peripheral tissues could significantly improve long-term biointegration of these devices (Adly et al., 2018; Minev et al., 2015). Additionally, the integration of non-planar electrode arrays that provide improved electrical access to the nerve – either by increasing efficacy of contact with the epineurim (Robinson et al., 2012; Shmoel et al., 2016) or by penetrating into the intraneural space (Angle et al., 2015; Gillis et al., 2018; Vitale et al., 2015) – may allow for greater recording and stimulating specificity with low risk of increased host tissue trauma (Salatino et al., 2017). And finally, we showed that current-steering-based stimulation can achieve a high degree of modulatory specificity within the small geometry of the nanoclip, but our results also highlight the challenge of identifying multi-channel stimulation parameters that will produce and maintain a specific physiological or end effector state (Cassar et al., 2017; Grill et al., 1991; Veraart et al., 1993). Thus, full realization of multi-channel neuromodulation for precise control of PNS activity will require closed-loop, unsupervised methods for mapping stimulation patterns to functional outputs such that robust control of end state is possible. These technologies are active areas of investigation by us and others; taken together, these developments would open the way for the realization of viable clinical devices for long-term precision interfacing in the periphery.

## METHODS

### Fabrication of the nanoclip nerve interface

The nanoclip interface for chronic peripheral nerve mapping and control used fabrication procedures and design features similar to our recent report (Lissandrello et al., 2017). Key steps in the fabrication of the device are overviewed in Figure S1 with device dimensions (Figure S2) as follows: total thin-film device length, 40 mm; neck width, 250 μm electrode array width, 420 μm; electrode array thickness, 12 μm; longitudinal electrode pitch, 80 μm; transverse electrode pitch, 45 μm; gold interconnect line width, 10 μm; number of electrode sites, 6; overall dimensions of nanoclip, LxWxH = 300×300×400 μm. The key fabrication steps (recounted in Figure S1) are as follows: (i) thin-film, multi-site electrode was fabricated using polyimide thin-film microfabrication techniques (HD Microsystems PI-2610; Smania, S.R.L.) and subsequently wire-bonded to a printed circuit board to ease connection to test equipment; (ii) the electrode was mounted on a thin optical glass substrate (24×60 mm, #0-thickness cover glass, Gold Seal) with 150 μm-thick double-sided acrylic tape (Tape #9713, 3M), and a drop (∼0.5-2 μL) of liquid acrylic photoresist (IP-Dip, Nanoscribe, GmbH) was deposited over and beneath the recording array; (iii) the nanoclip base (designed with Solidworks2016, Dassault Systèmes) was printed through the optical glass substrate using a two-photon-polymerization-based, dip-in resonant direct laser writing (rDWL) process (Pearre et al., 2018); (iv) the glass substrate and sample were inverted, and the top half of the nanoclip was printed with the rDWL process; (v) the photoresist was developed by submerging the glass substrate, electrode, and nanoclip in propylene glycol methyl ether acetate (PGMEA; Sigma Aldrich) for 20 min, and the whole device rinsed in methoxy-nonafluorobutane (Novec 7100; 3M) to remove trace PGMEA residue; (vi) a three-electrode cell comprising the gold pads as working electrodes, an Ag|AgCl reference electrode, and a large surface area platinum counter electrode was used with an iridium tetrachloride-oxalic acid plating solution to deposit EIROF via cyclic voltammetry (CV) (20-60 cycles at 50 mV/s sweep rate between −0.05 and 0.575 V) (Cogan, 2008; Meyer et al., 2001); (vii) the electrodes were then rinsed with de-ionized water and characterized by CV and electrochemical impedance spectroscopy (EIS) in 1x phosphate buffered saline (PBS) (see Figure S3; electrodes were plated to impedance, Z = 10-20 kΩ at 1kHz and charge storage capacity, CSC = 25-40 mC/cm^2^). To confirm integrity of fabrication, a subset of finished devices were sputter-coated with gold (3 cm distance to gold target; 1 min sputtering at 0.05 mbar and 20 mA; Sputter Coater 108) and imaged via SEM micrographs (6 mm working distance with the secondary electron sensor, 3 kV accelerating voltage, 30 μm aperture; Zeiss Supra 55VP; see Figure 1c). Though all tested devices were fabricated using the previously described rDLW system (Pearre et al., 2018), initial prototyping of the nanoclip structure was performed with a Nanoscribe Photonic Professional GT (Nanoscribe, GmbH).

### Vertebrate Animal Subjects

Adult male zebra finches (*Taeniopygia guttata*; 90+ days after hatch, n = 22 birds) were obtained from the Boston University breeding facility and housed on a 13:11 hr light/dark cycle in individual sound-attenuating chambers with food and water provided *ad libitum*. The care and experimental manipulation of the animals were performed in accordance with the guidelines of the National Institutes of Health and were reviewed and approved by the Boston University Institutional Animal Care and Use Committee. Because the behavioral effects of our interventions could not be pre-specified prior to the experiments, we chose sample sizes that would allow for identification of outliers and for validation of experimental reproducibility. No animals were excluded from experiments post-hoc. The investigators were not blinded to allocation of animals during experiments and outcome assessment, unless otherwise stated.

### Surgical procedures

All surgical procedures were performed under isoflurane anesthesia (1-4% dissolved in oxygen), with peri-operative analgesia (1% Lidocaine, SC) and anti-inflammatory (1% Meloxicam, IM) regimens. All nerve implants reported in this study targeted the avian hypoglossal cranial nerve, the tracheosyringeal nerve (nXIIts). The nXIIts, which runs along the length of the songbird trachea and terminates at the syrinx, has a diameter of approximately 150 μm and is composed of both afferent and efferent fibers (Lissandrello et al., 2017). At all experiment end-points, animals were given an overdose injection of sodium pentobarbital (250 mg/kg Euthasol, IC).

#### Acute preparation (n = 6 birds)

An anesthesia mask was placed over the bird’s head and the animal placed in a supine position with a small pillow was placed beneath the neck for support. Feathers were removed from the lower head, neck and upper chest, and betadine antiseptic solution (5% povidone-iodine) and ethanol (70%) were successively applied to prepare the incision site. A 20–25 mm incision was made at the base of the neck, and the tissue blunt dissected to expose the trachea. Sutures were placed in the skin of the lateral edge of the incision and retracted from the body to expose the implant site. Connective tissue surrounding the nerve was blunt dissected away, and two sections of the nXIIts (each 3–4 mm and approximately 15 mm apart) were isolated from the trachea. A nanoclip interface was implanted at the rostral location. For the recording experiments reported in Figures 3, a bipolar silver hook stimulating electrode was placed at the caudal location; for the stimulating experiments reported in Figures 5a-b, a second nanoclip interface was implanted at the caudal location. For all recording experiments, a platinum ground wire (0.003 in dia., Teflon coating; AM-Systems) was sutured to the inside of the skin and away from the neck muscles. Tissue dehydration during the procedure was minimized with generous application of PBS to the nerve and surrounding tissues. At the conclusion of the experiment, the animals were sacrificed and the devices recovered. Individual acute recording and stimulating experiments lasted 2–3 hr in total.

#### Fictive singing preparation (n = 6 birds)

The bird was prepared for surgery as above, with additional preparation of the skin at the abdomen at the caudal end of the sternum. A single 3-4 mm section of nXIIts was isolated from the trachea and implanted with a nanoclip interface, and the skin incision closed (around the protruding electrode neck) with 2-3 simple interrupted sutures. With the bird in lateral recumbency, a 2 mm silicone cannula was placed in the abdominal air sac (Brown and Pilny, 2006), secured to the skin with finger-trap suturing, and the air sac-cannula interface sealed with Kwik-Kast (WPI). The bird was wrapped in elastic nylon mesh and isoflurane delivery switched from the mask to the air sac cannula for the remainder of the experiment. At the conclusion of the experiment, the animals were sacrificed and the devices recovered. Individual fictive singing experiments lasted 3–5 hr in total.

#### Chronic implant for nerve function assessment (n = 9 birds)

The bird was prepared for surgery and trachea exposed via blunt dissection as in the *Acute Preparation*. Sections of the nXIIts (each 3–4 mm) were isolated bilaterally from the trachea. Each bird received one of the following manipulations: (i) bilateral implant of (dummy) nanoclip nerve interface (n = 3 birds); (ii) bilateral sham implant (n = 3 birds); (iii) bilateral nerve crush (transient ∼1 N force applied to nXIIts with 2.5 mm wide flat forceps; n = 3 birds). Anti-inflammatory splash block (∼250 μL 1% Meloxicam) was applied directly to implant sites, and incisions were closed with sutures. All birds exhibited normal rates of singing within 2 days of surgery.

#### Chronic implant preparation (n = 5 birds)

The bird was prepared for surgery as in the *Acute Preparation* and placed in a stereotax. A sagittal incision was made along the top of the head and the tissue retracted. Four to six stainless steel anchor pins (26002-10, Fine Science Tools) were threaded between layers of the skull, and a head cap was made from dental acrylic. The device connector was secured to the head cap with additional dental acrylic, and the nanoclip and a platinum reference wire (0.003 in dia., Teflon coated; AM-Systems) were trocared beneath the skin to the neck. The animal was then removed from the stereotax and placed in supine position with neck support. A 10-15 mm incision was made at the base of the neck, and the trachea and nXIIts isolated as described above. The nanoclip interface was implanted on the nerve, and the reference wire secured to the underside of the skin. Anti-inflammatory splash block (∼250 μL 1% Meloxicam) was applied directly to the implant site, and incisions were closed with sutures. All birds exhibited normal rates of singing within 2 days of surgery.

### In vivo Electrophysiology

All experiments were implemented and controlled using custom LabVIEW (National Instruments) and MATLAB (MathWorks) software applications.

#### Acute electrophysiology

Acute electrophysiological data were recorded using nanoclip interfaces with a RZ5 BioAmp Processor and an RA16PA Medusa Preamplifier (Tucker-Davis Technologies). Neural signals were digitized at 24.4 kHz and 16-bit depth and were Bessel bandpass filtered (1 Hz – 10 kHz, zero-phase). Stimulation currents were delivered – through either bipolar silver hook electrodes or a nanoclip interface – using a PlexStim programmable stimulator (Plexon). For all acute electrophysiology experiments, current pulses were biphasic, 200 μs/phase in duration, delivered at 1 Hz, and varied in amplitudes from −200 – 200 μA. By convention, positive current amplitudes are cathodic; negative amplitudes anodic.

#### Fictive singing

1-1.5% isoflurane in dissolved oxygen, warmed and humidified to near-physiological levels (35C, 80% humidity) by bubbler cascade to minimize respiratory tissue damage and prolong work time, was delivered by an air sac cannula (Figure 5d). Flow rates were adjusted to maintain a stable anesthesia plane while eliminating broadband noise from forced gas flow over the passive syringeal labia (typically 100-200 mL/min at 0.5-1.5 kPa). Fictive vocalizations were recorded with an omni-directional condenser microphone (AT-803, Audiotechnica) placed 5-10 cm from the bird’s open beak, amplified and bandpass filtered (10×, 0.1 – 8 kHz; Ultragain Pro MIC2200, Behringer), and digitized by data acquisition boards (PCIe-6212, National Instruments) and sound-triggered LabVIEW software at 44.15 kHz and 16-bit depth. Current-controlled stimulation was delivered through a nanoclip interface using a programmable stimulator (PlexStim, Plexon) with a custom MATLAB interface. 1 kHz bursts of 100 biphasic pulses, 200 μs/phase in duration, delivered at 0.25 Hz, and varying in amplitudes from −200 – 200 μA were delivered at each active stimulation site (of up to 6 in total). The current-steering parameters used here consisted of spatially distinct stimulation patterns (all current steering patterns tested are listed in Figure S4).

#### Chronic electrophysiology

All birds were recorded continuously using sound-triggered software as above, generating a complete record of vocalizations and nerve activity for the experiment. Neural recordings were acquired with an RHD 2000 system with a 16-channel unipolar input headstage (Intan Technologies), amplified, and bandpass filtered (0.3 – 15 kHz). Singing-related nerve activity was recorded from up to six sites on the nXIIts in n = 5 birds; recordings were made for >30 days in n = 3 birds. Across birds, we found high correlation between simultaneous recordings made at adjacent recording sites; we report on data collected at the most stable recording site in each bird, though we note that the trends were similar across all channels.

### Data analysis

All song and electrophysiology data analysis was performed off-line using custom-written MATLAB software.

#### Stimulation evoked responses

Stimulation-evoked responses were recorded from the right-side nXIIts. We sampled activity ∼5 ms before and up to 25 ms after stimulation onset and used the onset of the stimulation artifact (Figures 3b,d and 5b,c at 0 ms) to temporally align individual trial responses. Absolute response amplitudes were observed and quantified in a stimulation response window 0.75 – 4 ms after stimulation onset—a latency consistent with estimated nerve conduction velocities for 5 – 8 *μ*m diameter myelinated axons (i.e. 4 – 24 m/s; (Andrew and Part, 1972; Gillis et al., 2018). Signal-to-noise ratios (SNRs) were calculated from mean recordings (n = 20 trials) made in the stimulation response window (i.e., signal) and 10.75 – 14 ms after stimulation (i.e., noise). We considered an evoked response to be detected if the SNR within the signal response window exceeded a 90% confidence interval calculated by bootstrap (i.e., resampling with replacement the signal and noise intervals over n = 10000 trials) (Parks et al., 2016). Figures 3b and 5b show individual stimulation trials from single experimental sessions. Data points in Figure 3c show mean response over n = 20 trials for each bird; lines indicate individual devices/birds. Figure 3d shows the mean (solid line) and standard error (shaded region) across assayed devices.

#### Syllable segmentation and annotation

Raw audio recordings were segmented into syllables as previously described (Ali et al., 2013). Briefly, spectrograms were calculated for all prospective syllables, and a neural network (5,000 input layer, 100 hidden layer, 3–10 output layer neurons) was trained to identify syllable types using a test data set created manually by visual inspection of song spectrograms. Accuracy of the automated annotation was verified by visual inspection of a subset of syllable spectrograms.

#### Syllable feature quantification

Motif syllables were characterized by their pitch, frequency modulation, amplitude modulation, Wiener entropy, and sound envelope – robust acoustic features that are tightly controlled in adult zebra finch song (Ravbar et al., 2012; Tchernichovski et al., 2000). Each feature was calculated was calculated for 10 ms time windows, advancing in steps of 1 ms, such that an estimate was computed for every millisecond.

#### Song similarity

Quantification of acoustic similarity between syllable renditions produced over days was performed using a modified version of a previously described method for calculating acoustic similarity (Tchernichovski et al., 2000). Briefly, pairwise millisecond-by-millisecond comparisons of the similarity of acoustic features of identified syllables were made in five-dimensional space. The median distance between points was converted into a P-value based on the cumulative distribution of distances calculated between 20 unrelated. Thus, on this scale a similarity score of 1 means that syllables are acoustically identical, while a score of 0 indicates that the sounds are as different as two unrelated syllables. Quantification for each experimental group was calculated as the mean across syllables within a bird (100 renditions each of 3-6 unique syllable types) and then the mean over birds within a treatment group (n = 3 birds per group). Figure 2b shows the mean (solid line) and standard deviation (shaded region) for each treatment group.

#### t-SNE embedding of fictive vocalizations

For each bird (n = 6), fictive vocalizations produced from ∼20 trials of up to 24 stimulation patterns were included in this analysis. Audio recordings of each fictive vocalization, starting at the onset of stimulation and lasting for 200 ms, were converted into spectrograms (5 ms window, 1 ms advance, 512-point nfft). Spectrogram rows corresponding to frequencies above 8 kHz were discarded, and the remaining matrices (200 time steps by 91 frequency bins for each vocalization) were transformed into 18,200-dimension vectors. The dataset dimensionality was reduced (to 50) with principal components analysis, and this data was subsequently embedded in two-dimensional space using t-distributed stochastic neighbor embedding (t-SNE) (van der Maaten and Hinton, 2008) with distances calculated in Euclidean space and a perplexity of 35 (Figure 5f). Pairwise distances between t-SNE embedded datapoints were calculated in Euclidean space (Figure 5g).

#### Alignment of the neural recordings to song

A dynamic time warping (DTW) algorithm was used to align individual song motifs to a common template as previously described (Ali et al., 2013). The warping path derived from this alignment was then applied to the corresponding common mode subtracted and bandpass filtered nXIIts voltage recordings (0.3-6 kHz, zero-phase, 2-pole Butterworth) with no premotor time-shifting. The aligned neural traces were squared (to calculate signal envelope) and smoothed (Figure 5b: 20 ms boxcar window; all other analysis: 5 ms boxcar window, 1 ms advance).

#### Activity stability correlation

The stability of recorded nXIIts temporal dynamics was calculated as the Pearson’s correlation between the aligned neural signal envelope (averaged over 25 consecutive motifs) on the first day of recording with the same at later timepoints. The day 1 data point in Figure 4f denotes the correlation between the mean signal envelopes for two consecutive blocks of 25 motifs recorded on the first day. The running correlation (Figure 4d) shows Pearson’s correlation between the mean activity envelope of 25 motifs on the first day of recording and the mean of signal envelopes in a sliding window (width: 25; advance: 1).

#### Mean signal envelope amplitude

The mean signal envelope amplitude was calculated per motif and averaged over the 25-motif windows as described for the activity correlation (above). For analyses pooled across birds, mean envelope amplitude was normalized to the mean on the first day of recording.

### Statistical analysis

All statistics on data pooled across animals is reported in the main text as mean ± SD and depicted in figure error bars as mean ± SD, unless otherwise noted. Figure starring schema: *p<0.05, **p<0.01, ***p<0.001. Where appropriate, distributions passed tests for normality (Kolmogorov-Smirnov), equal variance (Levene), and/or sphericity (Mauchly), unless otherwise noted. Multiple comparison corrected tests were used where justified. Statistical tests for specific experiments were performed as described below.

*Figure 2b:* Comparison of pre- and post-surgical song acoustic structure following bilateral nanoclip implant (n = 3 birds), nerve crush (n = 3 birds), and sham controls (n = 3 birds). A two-tailed, paired t-test revealed significant differences in acoustic structure immediately after nerve crush (P = 0.02), but not for nanoclip or sham implants (P = 0.07 and P = 0.53, respectively). In addition, two-tailed unpaired t-tests showed significant differences between nerve crush and sham controls at all post-procedure time points tested (i.e., through the eighth day: P < 0.01); no significant differences were found between nanoclip implants and sham controls at any time point (P > 0.22).

*Figure 3e:* Comparison of stimulation evoked response amplitudes before and after lidocaine/saline application in n = 3 birds. Mauchley’s test indicated a violation of sphericity (W = 0, P = 0), and a Huynh-Feldt degree of freedom correction was applied. Subsequent repeated-measures ANOVA revealed significant differences between the treatments (F_(1.11,2.21)_ = 189.02, P = 0.003). Post-hoc comparisons using Dunnett’s test showed significant differences between saline (control) and lidocaine application (P = 6 × 10^−6^); no other condition significantly differed from control (P > 0.15).

*Figure 4d:* Comparison of nXIIts dynamics over 30 days of continuous recording in n = 3 birds. A two-tailed, paired t-test revealed no significant differences in correlation between recording Day 1 and Days 10, 20, and 30 (P > 0.05).

*Figure 4e:* Comparison of normalized nXIIts activity over 30 days of continuous recording in n = 3 birds. A two-tailed, paired t-test revealed no significant differences in mean activity envelope amplitude between recording Day 1 and Days 10, 20, and 30 (P > 0.06).

*Figure 5g:* Mean pairwise distance between t-SNE embedded fictive vocalizations within and across multi-channel stimulation patterns in n = 6 birds. A two-tailed, paired t-test revealed significantly greater mean distances across different stimulation patterns than within repeated application of the same pattern (P = 0.002).

### Immunohistology

At the end of the experiments, birds were anesthetized with sodium pentobarbital (Euthasol, IC) and subsequently transcardially perfused with cold PBS, followed by fixation with 4% paraformaldehyde (PFA) in PBS. The trachea and tracheosyringeal nerves (with nanoclip in place) were dissected out and post-fixed in 4% PFA overnight, then cryo-protected in 10% sucrose for an additional night. The tissue was flash-frozen in cutting medium (Tissue-Tek, OCT Compound; Sakura Finetek), and axial sections (35 µm) were made on a Cryostat (Leica) through both the tissue and the nanoclip. Slices were blocked (1% BSA, 0.3% Triton in PBS) at room temperature and incubated with anti-neurofilament (Chicken, 1:1000, Developmental Studies Hybridoma Bank, 3A10) and anti-NeuN (Mouse, Millipore, MAB377) primary antibodies in blocking buffer for 48 hrs at 4°C. After washing, slices were incubated with anti-Chicken-Alexa 568 (goat, 1:1000, Life Technologies, A-11041) and anti-Mouse-Alexa 647 (goat, 1:1000, Life Technologies, A-31625) over night at 4°C. Slices were mounted and imaged using an Olympus Fluoview FV10i confocal microscope for high resolution images.

## Code and data availability

All custom-written code is accessible in an online repository; data and device design files are available upon request.

## Acknowledgements

The authors thank Silvia Bossi for fabrication of the thin-film array at Smania, S.R.L. We also thank Ian Davison, Benjamin Scott, Jeff Gavornik, and members of the Gardner Lab for helpful comments on drafts of this manuscript. This research was supported by the NIH (R01NS089679) and a sponsored research agreement with GlaxoSmithKline. The nanoclip has been described in a patent filing (US Patent PR66142P, US Serial No. 62/367,975).

## Author Contributions

TMO, CM, AEW, and TJG conceptualized the project, designed the device, and developed the fabrication method with input from BJH and DJC. TMO, BL, and KG performed surgical procedures and in vivo experiments. TMO, KG, and JG analyzed experimental data. LD performed SEM sample preparation and imaging. DS performed immunohistology, biological sample imaging, and animal husbandry. TMO drafted the manuscript with input from all other authors.

## Supplementary Figures

**Figure S1.**
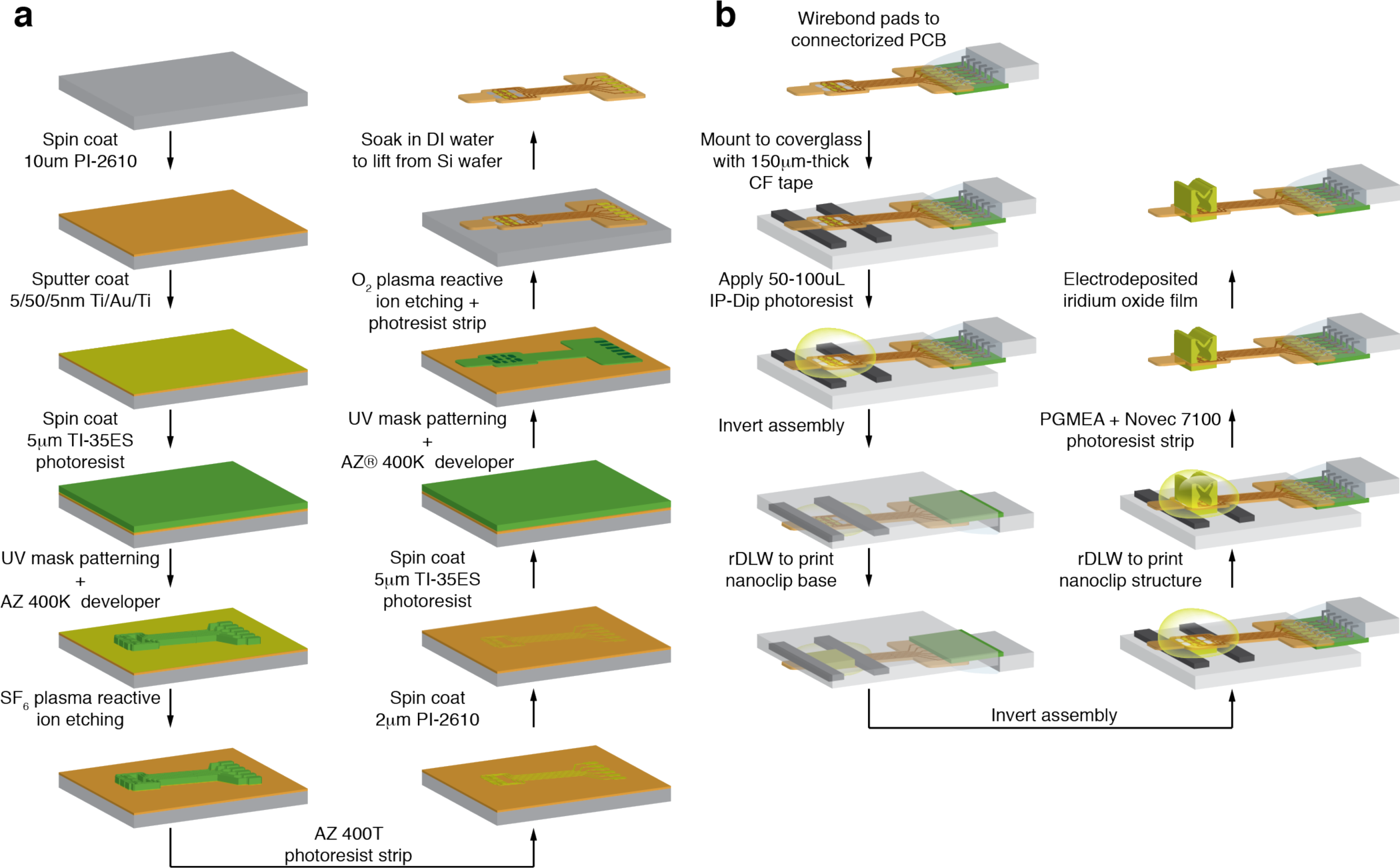
Nanoclip interface fabrication process. **(a)** Polyimide thin-film electrode fabrication process. (Custom thin-film microfabrication performed by Smania, Pisa, Italy). **(b)** Thin-film electrode and nanoclip integration process.

**Figure S2.**
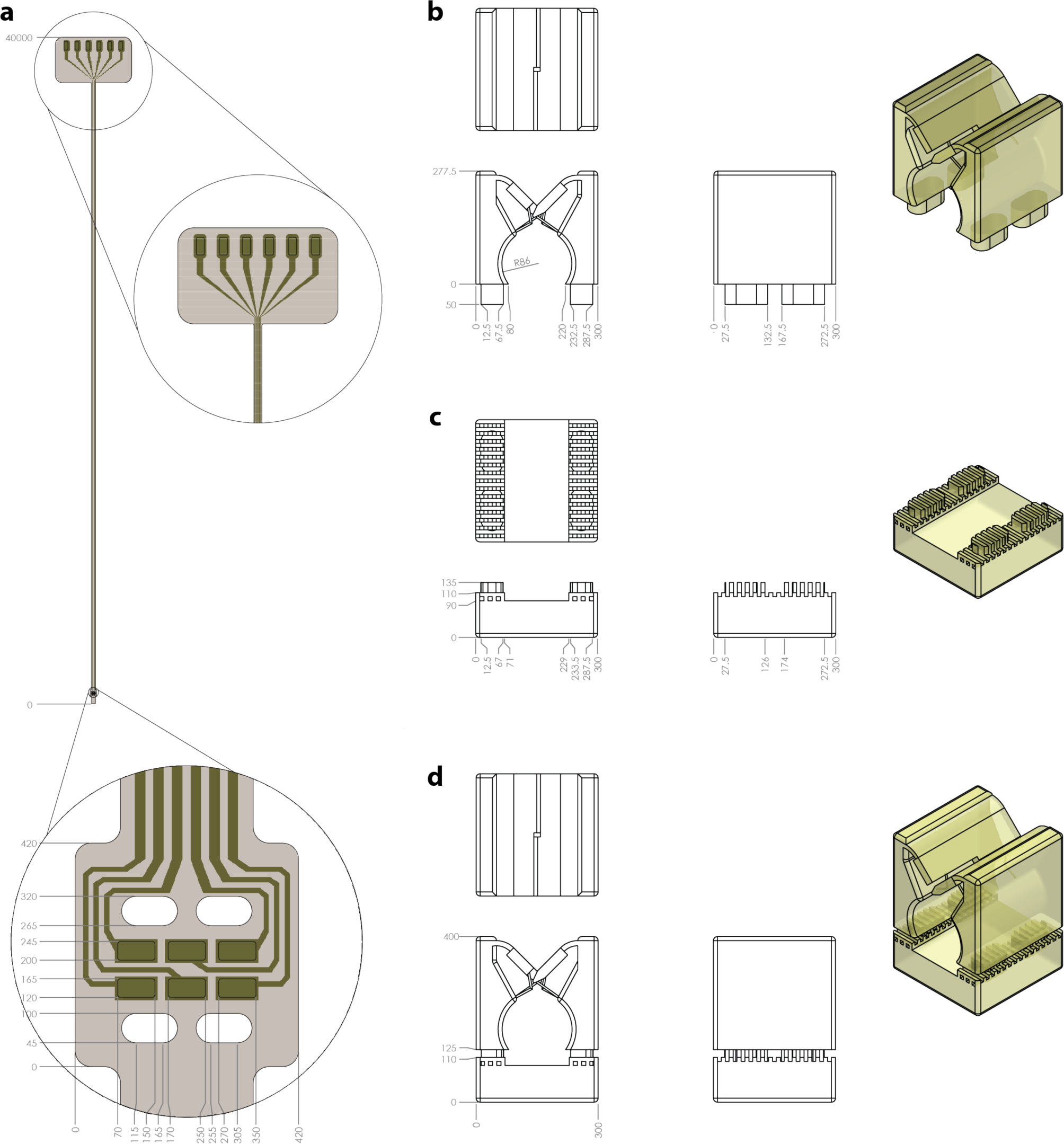
Design and geometry of the nanoclip interface components. **(a)** Diagram and dimensions of the polyimide thin-film electrode array. Top inset: Bottom inset: electrode array and through-vias. **(b-d)** Diagram and dimensions of the 3D-printed nanoclip: **(b)** upper structure comprising the interlocking trap doors and nerve cavity; **(c)** base structure with interleaved tenon; **(d)** as-fabricated integration of the nanoclip upper and base. All dimensions in microns.

**Figure S3.**
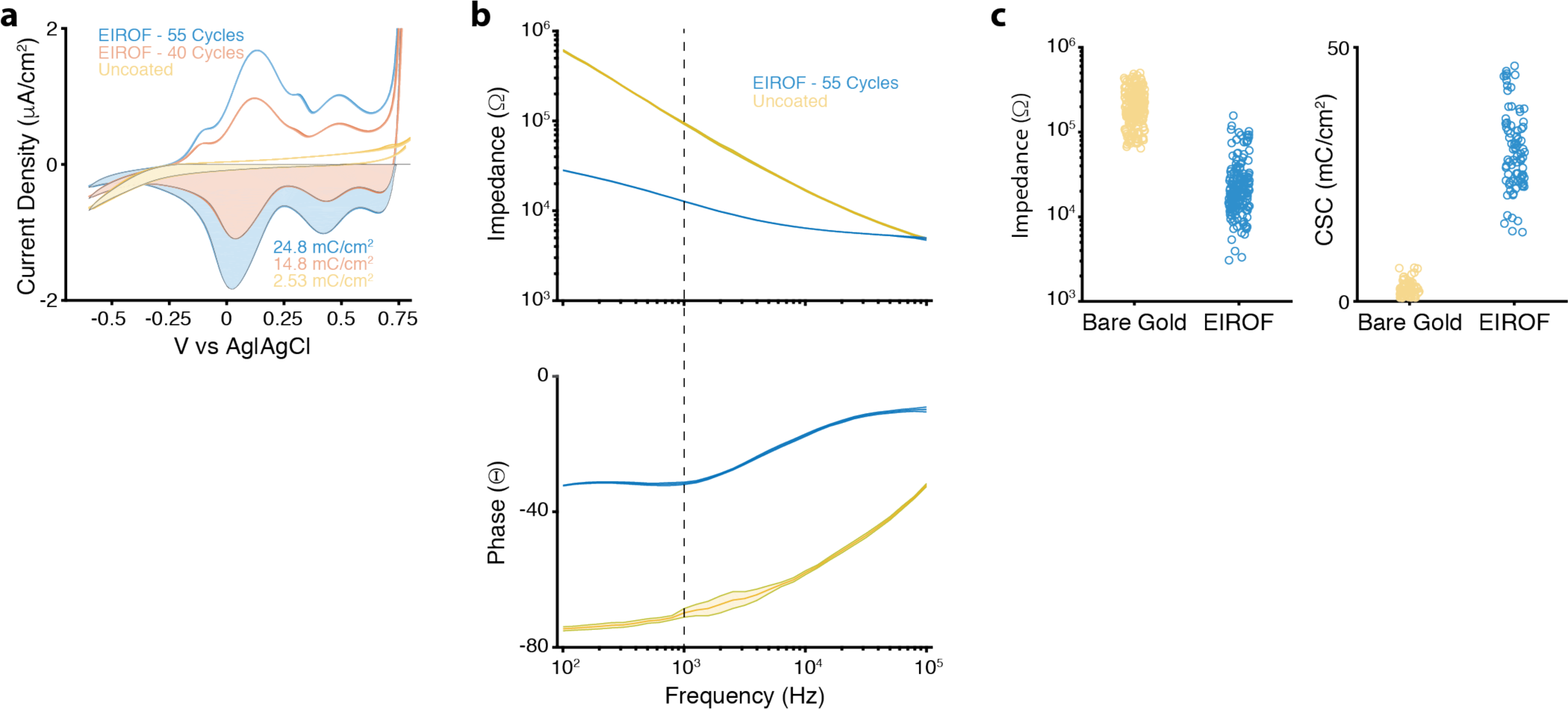
Electrodeposition of iridium oxide films (EIROF) to condition electrodes for neural recording and stimulation. **(a)** Representative cyclic voltammetry curves measured in PBS for a gold electrode before and after EIROF deposition. Shaded region shows time-integral of the negative current used to calculate charge storage capacity (CSC) as in (**c**) (Cogan, 2008). Yellow: uncoated gold. Red: 40 deposition cycles. Blue: 55 cycles. **(b)** Representative electrode impedance spectroscopy measured in PBS for a gold electrode before and after EIROF deposition. Top: Impedance versus frequency. Bottom: Phase versus frequency. Dashed line shows nominal values at 1 kHz as in (**c**). EIS Color as in (**a**). **(c)** Summary plots for impedance at 1 kHz (left) and CSC (right) before (yellow) and after (blue) IROF deposition (n = 210 pads in 35 devices and n = 90 pads in 15 devices, respectively). Data points correspond to individual electrode pads.

**Figure S4.**
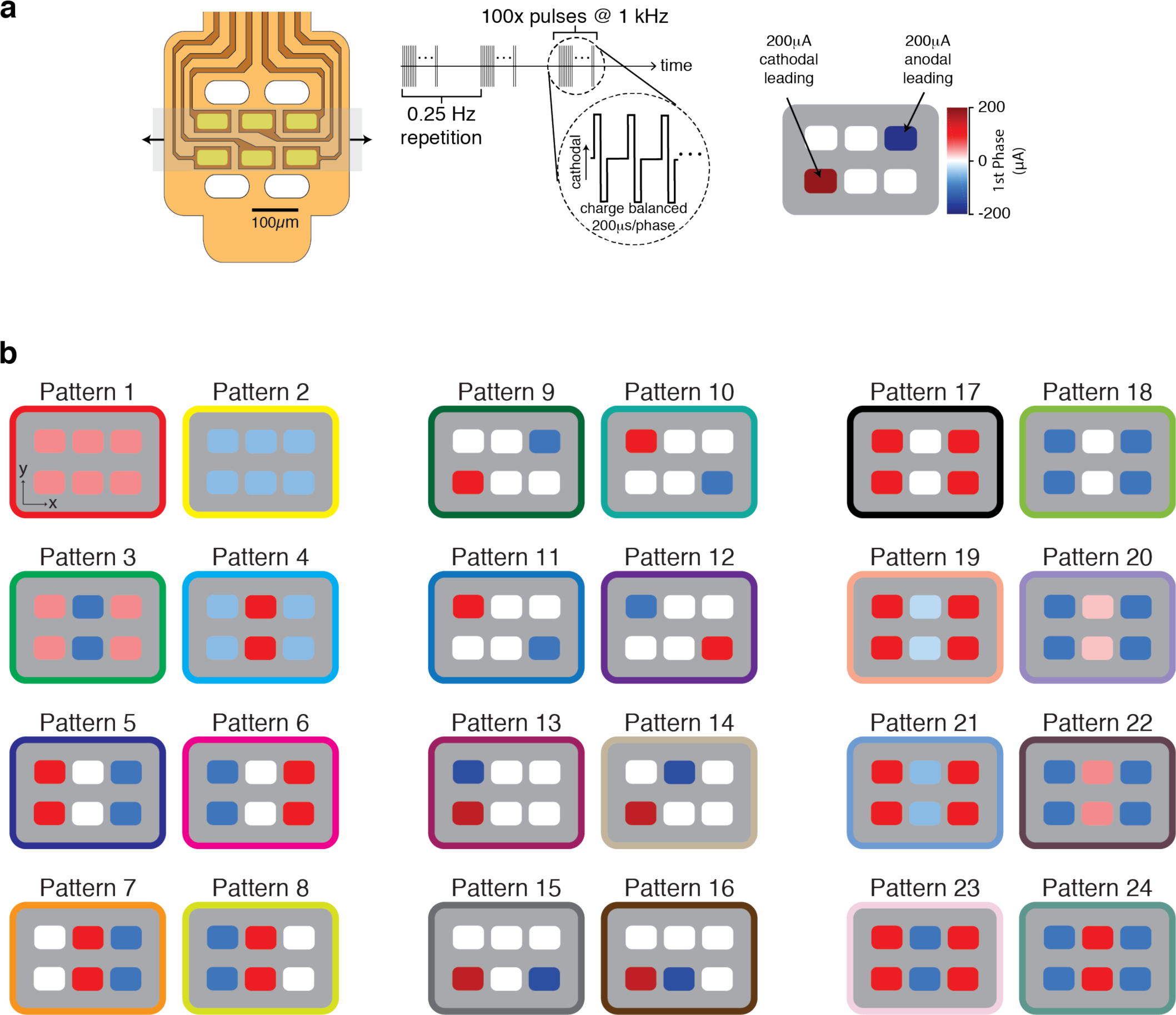
Current steering patterns for fictive singing experiments. **(a)** Schema for multi-channel stimulation patterns for fictive-singing experiments. (Left) Diagram of the six-channel electrode array used in all experiments. The shaded gray region shows the location of a retained nerve; the arrows indicate the nerve axis. (Right) 100 biphasic stimulating pulses, 200 μs/phase at −200-200 μA, were delivered at 1 kHz, repeated at 0.25 Hz. In the pattern schema, the color of each pad indicates the magnitude and direction of the leading phase of the stimulation pulse. **(b)** All current-steering stimulation patterns. The electrode pad color is as in (**a**). The color of the border indicates the stimulation patterns employed during the experiment reported in Figure 5e.

